# Disrupted sleep-wake regulation in the MCI-Park mouse model of Parkinson’s disease

**DOI:** 10.1101/2023.11.09.566376

**Authors:** K.C. Summa, P. Jiang, P. González-Rodríguez, X. Huang, X. Lin, M.H. Vitaterna, Y. Dan, D.J. Surmeier, F.W. Turek

## Abstract

Disrupted sleep has a profound adverse impact on lives of Parkinson’s disease (PD) patients and their caregivers. Sleep disturbances are exceedingly common in PD, with substantial heterogeneity in type, timing, and severity. Among the most common sleep-related symptoms reported by PD patients are insomnia, excessive daytime sleepiness, and sleep fragmentation, characterized by interruptions and decreased continuity of sleep. Alterations in brain wave activity, as measured on the electroencephalogram (EEG), also occur in PD, with changes in the pattern and relative contributions of different frequency bands of the EEG spectrum to overall EEG activity in different vigilance states consistently observed. The mechanisms underlying these PD-associated sleep-wake abnormalities are poorly understood, and they are ineffectively treated by conventional PD therapies. To help fill this gap in knowledge, a new progressive model of PD – the MCI-Park mouse – was studied. Near the transition to the parkinsonian state, these mice exhibited significantly altered sleep-wake regulation, including increased wakefulness, decreased non-rapid eye movement (NREM) sleep, increased sleep fragmentation, reduced rapid eye movement (REM) sleep, and altered EEG activity patterns. These sleep-wake abnormalities mirror those identified in PD patients.

Thus, this model may help elucidate the circuit mechanisms underlying sleep disruption in PD and identify targets for novel therapeutic approaches.

## Introduction

The defining features of Parkinson’s disease (PD) are the motor symptoms of bradykinesia, rigidity, and tremor, which are attributable to the degeneration of dopaminergic neurons in the substantia nigra pars compacta (SNc). Although levodopa-responsive motor disability is critical to the diagnosis of PD, these symptoms are frequently accompanied by a range of non-motor symptoms^1^, among the most common of which is disrupted sleep^2,3^. Sleep-wake disturbances are experienced by up to 80% of PD patients^3^. Recent systematic reviews have summarized what is known of sleep disturbances in PD patients^2–8^. The most prominent sleep-related symptoms include insomnia; excessive daytime sleepiness; and fragmentation^7^, which refers to interrupted sleep and can preclude accumulation of adequate sleep and dissipation of homeostatic sleep need. Sleep disturbances in PD are characterized by difficulty falling asleep and staying asleep (insomnia and fragmentation), as well as difficulty in maintaining daily sleep-wake cycles, with a reduction in the amplitude of day-night sleep rhythms as well as excessive daytime sleepiness and increased nocturnal awakenings, which are often reported in PD patients^5,6,8^. Interestingly, sleep abnormalities often precede PD motor symptoms^6^, and such cases are frequently associated with more severe motor symptoms, a treatment-refractory phenotype, and accelerated disease progression^7,9^.

Changes in brain wave activity, as measured by the electroencephalogram (EEG), have been demonstrated in PD patients, who typically exhibit an overall slowing of EEG activity as detected by relative increases in the proportion of lower frequency bands, such as delta and theta, and relative decreases in higher frequency bands, such as alpha, to the EEG spectral profile of different vigilance states^10–12^. Similar patterns of EEG slowing have been observed in several mouse models of PD^13–15^. Interestingly, particular EEG changes, such as theta power during wake, have been associated with cognitive performance, including in PD^10,16–19^. Together, these findings suggest the pathologic changes of PD are accompanied by specific EEG changes, which may therefore serve as important diagnostic, phenotypic, and prognostic disease activity markers.

Rapid eye movement (REM) sleep behavior disorder (RBD)^20^ has garnered particular interest due to its connection with PD. Most patients diagnosed with idiopathic RBD will ultimately progress to PD, Lewy body dementia (LBD), or multi-system atrophy (MSA) over time^2,21^. A recent meta-analysis including more than 17,000 PD patients estimated an overall pooled RBD prevalence of 46%^7^; however, a rigorous clinical RBD diagnosis incorporating the gold-standard of polysomnography is lacking in many studies, which makes it difficult to estimate prevalence reliably^7^. In addition to this diagnostic uncertainty, it seems likely that underlying pathogenic mechanisms may be distinctive given the lack of predictive specificity of RBD, with its associations to PD, LBD, and MSA^20^.

The pathophysiology driving sleep disturbances and sleep abnormalities in most PD patients is unclear. Alterations in brain circuitry responsible for PD symptoms and features have been studied primarily in animal models^22^. While some of these models exhibit sleep-wake abnormalities resembling those in PD patients^6,13–15,23–28^, no individual model faithfully captures the full spectrum of sleep disturbances present in PD^6^. Moreover, because these mouse models are not progressive, they do not allow for a rigorous dissection of the evolution and temporal dynamics of sleep disturbances as pathology unfolds.

Recently, a new mouse model of PD was introduced that was created using intersectional genetics to selectively disrupt mitochondrial complex I (MCI) function in dopaminergic neurons^29^. These mice, termed “MCI-Park mice,” exhibit a progressive levodopa-responsive form of parkinsonism. Moreover, the underlying pattern of neuropathology in these mice is strikingly similar to that inferred to be occurring in most PD patients^29^. To determine if MCI-Park mice also manifest PD-like disruption of sleep, both prodromal and parkinsonian mice were examined. These studies revealed that sleep abnormalities begin in young prodromal mice and progress. The patterns of sleep-wake abnormalities present in parkinsonian mice match the spectrum of those observed in PD patients: increased wakefulness, decreased non-REM (NREM) sleep, reduced amplitude of day-night sleep rhythms, increased fragmentation, slowing of EEG activity, and severely impaired REM sleep. These sleep abnormalities were seen in MCI-Park mice studied at two separate facilities, confirming their reproducibility in different environments. These studies suggest that the most common sleep abnormalities in PD patients may stem from the loss of dopaminergic neurons.

## Results

The experimental protocol with examples of sleep state epochs (i.e., wake, NREM sleep, and REM sleep) is depicted in Fig. 1. Younger animals (6-8 weeks of age) are in the prodromal state without overt evidence of motor dysfunction, whereas older animals (14-18 weeks of age) are in the symptomatic parkinsonian state and manifest characteristic motor deficits^29^. Sleep-wake behavior was recorded and analyzed as previously described^30,31^. Our previous comprehensive genetic analysis of sleep-wake traits in mice revealed that the traits cluster into distinct dimensions of related traits, or factors, such as state amounts, fragmentation, EEG power bands, and REM sleep)^31^. The results of the present studies are organized and presented according to these factors.

**Fig. 1:**
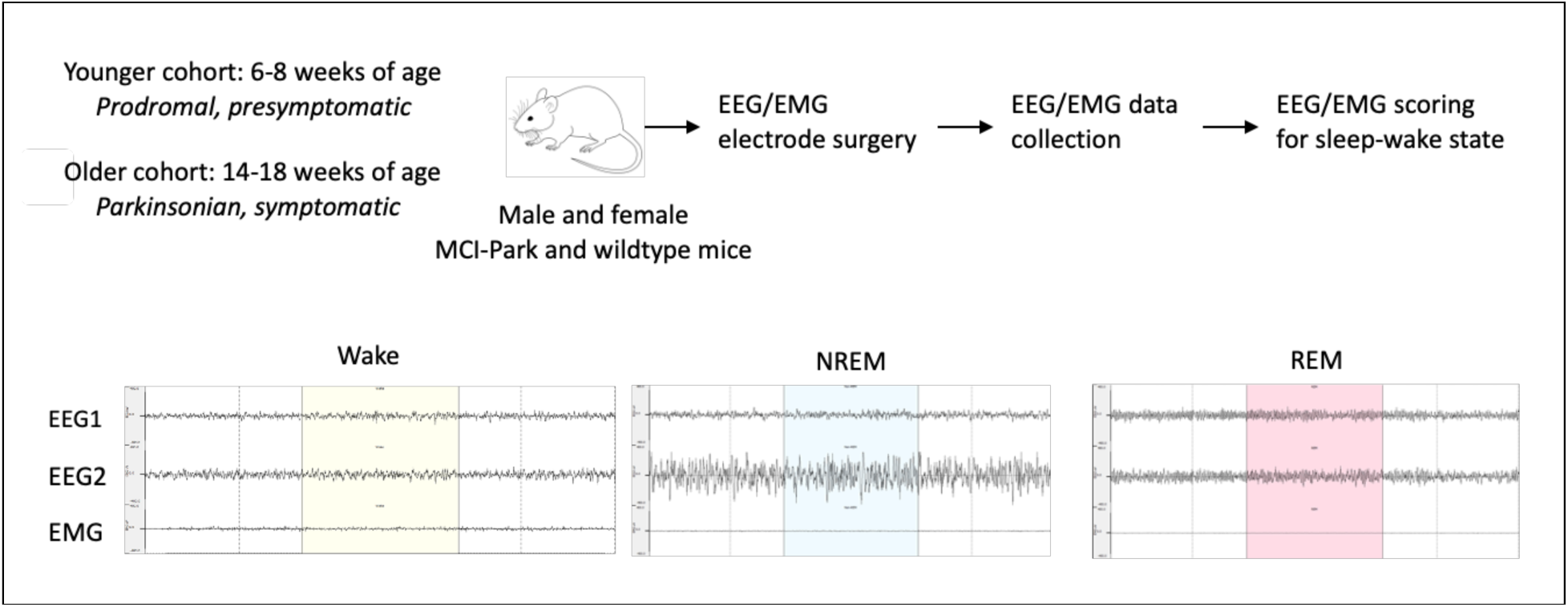
Experimental protocol. Male and female MCI-Park mice and wildtype littermates were obtained from a breeding colony maintained at Northwestern University. Mice ranging from approximately 37-57 days of age (younger cohort; presymptomatic or prodromal MCI-Park mice and age-matched wildtype littermates) and from approximately 88-121 days of age (older cohort; symptomatic parkinsonian MCI-Park mice and age- matched wildtype littermates) underwent electroencephalography (EEG) and electromyography (EMG) recording electrode implantation surgery and a minimum of 7 days of undisturbed recovery. EEG/EMG data were collected continuously and scored as wake, non-rapid eye movement (NREM) sleep, or rapid eye movement (REM) sleep as described. Representative 10-second example epochs for each state are depicted. Wake is characterized by low amplitude high-frequency EEG waves with variable EMG activity. NREM sleep is characterized by high-amplitude low-frequency EEG waves with absent EMG activity. REM sleep is characterized by low amplitude high-frequency EEG waves with absent EMG activity. Descriptions and definitions of each sleep-wake trait examined are provided in the Supplemental Material.

### MCI-Park mice had reduced sleep

The amount of time spent awake and in NREM sleep over 24 hours is depicted in Fig. 2 (REM sleep is presented separately below). Significant differences in wake time were detected between genotypes (*p<*0.001, F(1,65)=15.95; Fig. 2a), with MCI-Park mice exhibiting increased wakefulness. This was affected by age (genotype X age interaction *p*<0.001, F(1,65)=16.53), with older MCI-Park mice spending the most time awake. MCI-Park mice exhibited less total sleep (NREM sleep plus REM sleep) and NREM sleep over 24 hours (significant differences between genotypes for total sleep *p*<0.001, F(1,65)=15.97; and for NREM sleep *p*<0.01, F(1,65)=10.78; Fig. 2b). These differences were also impacted by age (genotype X age interaction *p*<0.001, F(1,65)=16.52 for total sleep, and *p*<0.001, F(1,65)=12.61 for NREM sleep), with older MCI-Park mice getting the least amount of NREM sleep. Older animals and MCI-Park mice had more NREM sleep as a proportion of total sleep (*p*<0.001, F(1.65)=27.26 for older *vs* younger mice and *p*<0.05, F(1,65)=4.25 for MCI-Park *vs* wildtype; Fig 2c). MCI-Park mice also exhibited a lower proportion of total sleep during the light phase of the light-dark cycle (*p*<0.05, F(1,65)=6.03; Fig 2d), indicating in a reduction in the amplitude of the day-night sleep-wake rhythm (i.e., MCI-Park mice sleep less during the light phase of the light-dark cycle, the “right” time of day for nocturnal animals to sleep).

**Fig. 2:**
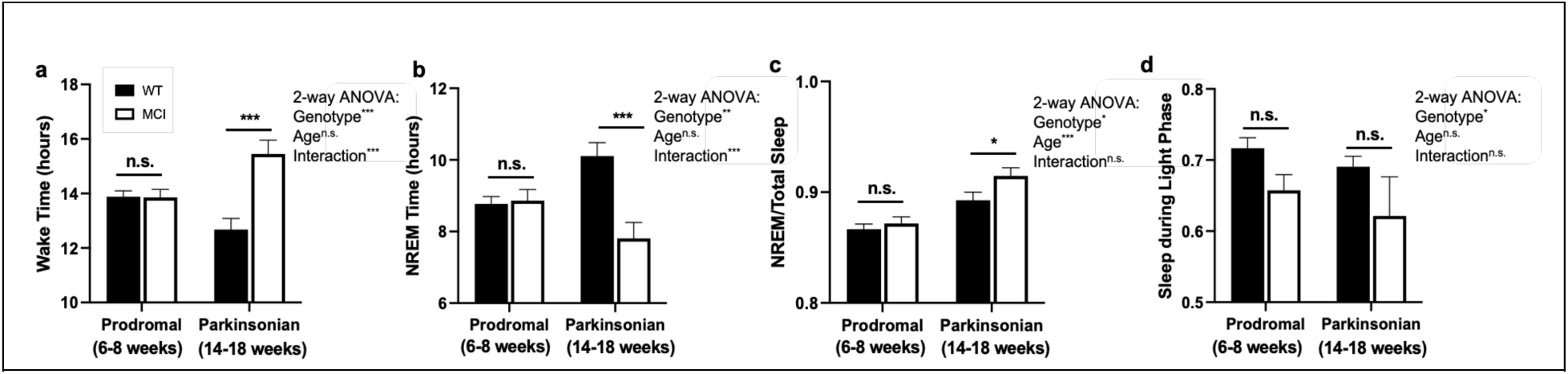
Wake and NREM sleep amounts and proportion of total sleep during the light phase. **a** – total time spent awake over 24 hours in wildtype (filled bars) and MCI-Park mice (open bars) at 6-8 weeks of age and 14-18 weeks of age. **b** – total time spent in NREM sleep over 24 hours in wildtype and MCI-Park mice at 6-8 weeks of age (filled bars) and 14-18 weeks of age (open bars). **c** – proportion of total sleep that is NREM sleep (NREM sleep amount/total sleep amount) in wildtype (filled bars) and MCI-Park (open bars) mice at 6-8 weeks of age (solid bars) and 14-18 weeks of age (open bars). **d** – proportion of total sleep (NREM plus REM sleep) during light phase of the light-dark cycle. *, *p*<0.05; **, *p*<0.01; ***, *p*<0.001. *N*=10-24 mice per genotype per age.

### MCI-Park mice had increased sleep fragmentation

Fragmentation occurs when sleep is interrupted. It limits the consolidation of sleep, it prevents accumulation of total sleep amount, and it impairs dissipation of sleep pressure that builds with increasing time awake, a key homeostatic mechanism of sleep regulation.

Fragmentation incorporates both the number and duration of bouts of different vigilance states: sleep can be fragmented due to changes in the total number of bouts of a state (i.e., more bouts that are less consolidated), changes in the duration of bouts of that sleep-wake state (i.e., shorter bouts), or to a combination. Significant differences in the number of wake bouts (Fig. 3a) were detected between genotypes (*p*<0.001, F(1,65)=21.64) and ages (*p*<0.001, F(1,65)=23.95), with MCI-Park and older mice having significantly more wake bouts than wildtype and younger mice, respectively. Differences in median wake bout duration were not observed between genotypes and ages (Fig. 3b). Significant differences in the number of NREM bouts (Fig. 3c) were also detected between genotypes (*p*<0.001, F(1,65)=15.03) and between ages (*p*<0.01, F(1,65)=15.68), with MCI-Park and older mice experiencing significantly more NREM bouts compared to wildtype and younger mice, respectively. Median NREM bout duration (Fig. 3b) was significantly different between genotypes (*p*<0.001, F(1,65)=3.181e-08) and ages (p<0.01, F(1,65)=5.739e-03), with MCI-Park and older mice having a significantly shorter median NREM bout duration compared to wildtype and younger mice. Taken together, MCI-Park and older mice exhibited significantly more bouts of wake and NREM sleep over the course of the day, with bout durations that were similar for wake and shorter for NREM sleep.

**Fig. 3:**
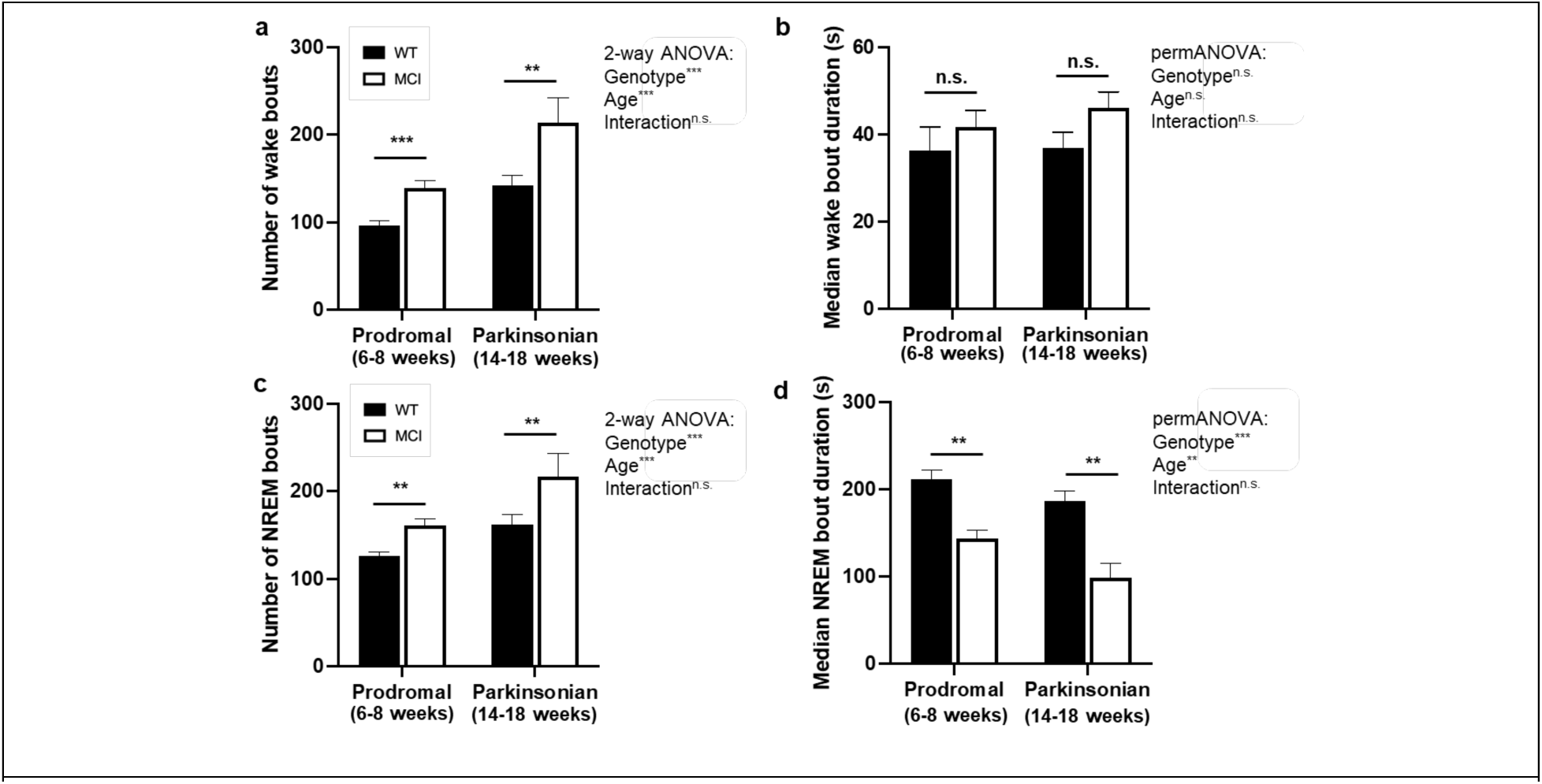
Wake and NREM bout number and median bout duration. **a** – number of wake bouts over 24 hours in wildtype (filled bars) and MCI-Park mice (open bars) at 6-8 weeks of age and 14-18 weeks of age. **b** – median wake bout duration in wildtype (filled bars) and MCI-Park (open bars) mice at 6-8 weeks of age and 14-18 weeks of age. *, *p*<0.05; **, *p*<0.01; ***, *p*<0.001. *N*=10-24 mice per genotype per age.

Additional measures of fragmentation include the traits known as “state shifts,” defined as the number of times that the observed sleep-wake epoch is different from the previous one, and “brief arousals,” defined as the number of single wake epochs occurring within the middle of a sleep bout^31^. Significant differences in state shifts (Fig. 4a) were detected between genotypes (*p*<0.01, F(1,65)=11.77) and ages (*p*<0.001, F(1,65)=17.90), with MCI-Park and older mice having more state shifts compared to wildtype and younger mice, respectively. A significant difference between genotypes was not observed for brief arousals (Fig. 4b), though there was a significant effect of age (*p*<0.001, F(1,65)=12.64), with the older mice exhibiting more brief arousals.

**Fig. 4:**
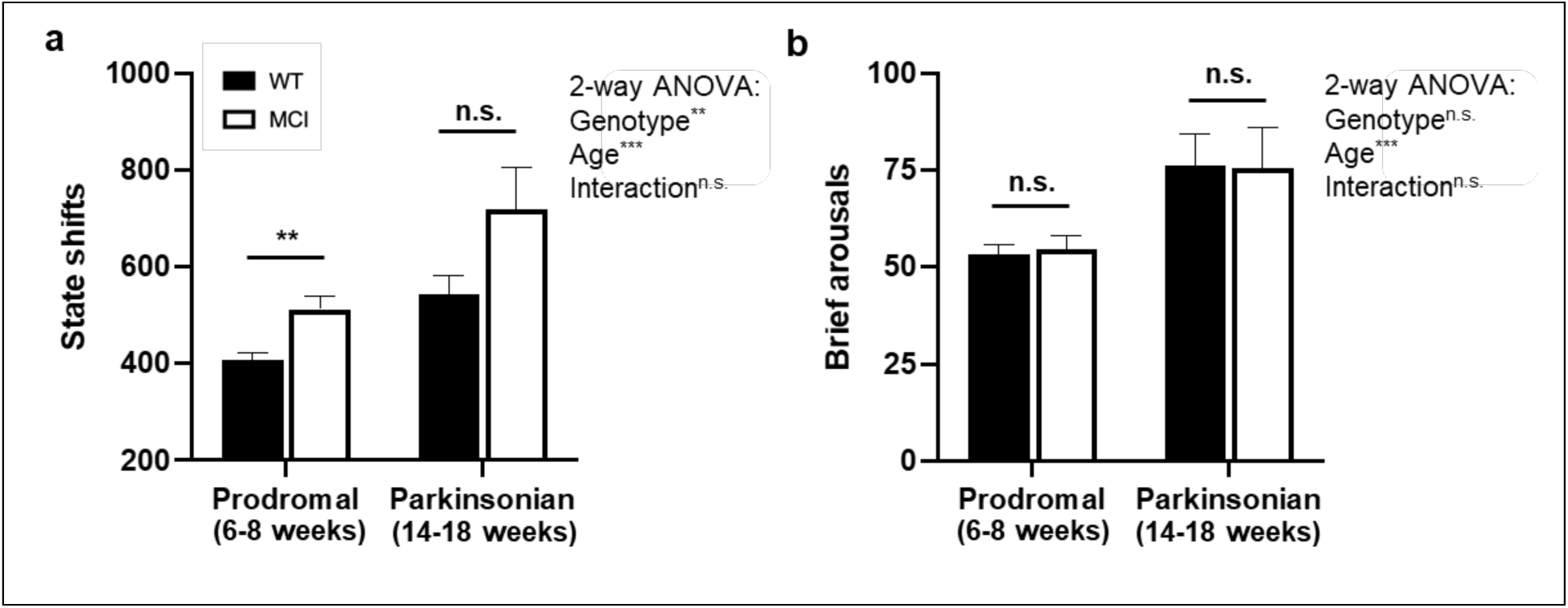
State shifts and brief arousals. **a** – number of state shifts over 24 hours in wildtype (filled bars) and MCI-Park (open bars) mice at 6-8 weeks of age and 14-18 weeks of age. **b** – number of brief arousals over 24 hours in wildtype (filled bars) and MCI-Park (open bars) mice at 6- 8 weeks of age and 14-18 weeks of age. *, *p*<0.05; **, *p*<0.01; ***, *p*<0.001. *N*=10-24 mice per genotype per age.

### MCI-Park mice had altered EEG spectra

The relative EEG power for each vigilance state (i.e., wake, NREM sleep, and REM sleep) for each frequency band (delta 0.5-4 Hz, theta 4-8 Hz, alpha 8-11 Hz, sigma 11-15 Hz, and beta 15-30 Hz) of the EEG spectrum for MCI-Park and wildtype mice in both the prodromal and parkisonian states is presented in Supplemental Table 1. This is calculated as the power for that frequency band divided by the total power (i.e., power of all frequency bands together) for that particular vigilance state and expressed as a percentage. Lower relative power indicates a smaller relative contribution of EEG activity within that frequency band to overall activity.

Conversely, higher relative power indicates more activity within that frequency band for the vigilance state. This spectral quantification of EEG activity profiles provides profiles of brain wave activity for the different vigilance states that may be useful to infer underlying neurologic and physiologic processes. A consistent finding in MCI-Park mice across each of the vigilance states is a significant decrease in the relative power in the alpha band (8-11 Hz) (Table 1). In NREM sleep, MCI-Park mice exhibited increased relative power in the delta band (0.5-4 Hz) (Table 1). There was also a significant effect of age, with the older mice demonstrating increased NREM delta power compared to younger animals. Taken together, in MCI-Park mice there is a shift towards slower EEG patterns, with a reduction in the relative contribution of the higher frequency (faster) alpha band to overall power, in conjunction with an increase in the relative contribution of the lower frequency (slower) delta band during NREM sleep in particular.

**Table 1:** EEG Power Bands. Relative EEG spectral power in the alpha (8-11 Hz) and delta (0.5-4 Hz) bands during wake, NREM sleep, and REM sleep in wildtype and MCI-Park mice. Relative power is calculated as the power of that particular frequency band divided by the overall power of the signal and is expressed as a percentage ± standard error. Data analyzed by generalized linear models (GLM) two-way analysis of variance (ANOVA). *, *p*<0.05; **, *p*<0.01; ***, *p*<0.001. *N*=10-23 mice per genotype per age.

| State | Power Band | WT Prodromal<br>(6-8 weeks) | WT Parkinsonian<br>(14-18 weeks) | MCI Prodromal<br>(6-8 weeks) | MCI Parkinsonian<br>(14-18 weeks) | Genotype | Age | Interaction |
|---|---|---|---|---|---|---|---|---|
| Wake | Alpha<br>(8-11 Hz) | 15.6±0.8% | 15.4±0.4% | 14.0±0.6% | 12.2±1.3% | **\*\*, p<0.01, F=8.0** | F=1.6, p=0.21 | F=1.1, p=0.30 |
| NREM | Delta<br>(0.5-4 Hz) | 39.3±1.5% | 42.5±2.1% | 46.8±1.5% | 49.1±2.0% | **\*, p<0.05, F=4.9** | **F=12.5, p<0.001** | F=1.6, p=0.24 |
| NREM | Alpha<br>(8-11 Hz) | 10.3±0.5% | 12.1±0.5% | 9.6±0.4% | 9.9±0.4% | **\*\*, p<0.01, F=7.3** | F=3.0, p=0.09 | F=1.5, p=0.23 |
| REM | Alpha<br>(8-11 Hz) | 16.7±0.7% | 19.9±0.8% | 15.0±0.6% | 14.4±1.5% | **\*\*\*, p<0.001, F=14.2** | F=1.4, p=0.24 | F=3.9, p=0.49 |

### MCI-Park mice had impaired REM sleep

Significant differences in the amount of REM sleep over 24 hours (Fig. 5a) were observed between genotypes (*p*<0.001, F(1,65)=14.07) and ages (*p*<0.001, F(1,65)=22.33), with MCI-Park and older mice exhibiting significantly less REM sleep compared to wildtype and younger mice. These differences were affected by a significant genotype X age interaction (*p*<0.01, F(1,65)=8.40), with older MCI-Park mice having the least REM sleep: over 40% less on average than the younger MCI-Park mice. The proportion of total sleep spend in REM was also reduced in MCI-Park compared to wildtype mice (*p*<0.05, F(1,65)=4.247) and in older compared to younger mice (*p*<0.001, F(1,65)=27.26), indicating that REM sleep contributes less to overall sleep amount in the MCI-Park and older animals. In addition, the proportion of REM sleep occurring during the light phase of the light-dark cycle was lower in MCI-Park compared to wildtype mice (*p*<0.05, F(1,65)=5.19), highlighting a reduction in the amplitude of the day-night rhythm of REM sleep in MCI-Park mice, as seen with NREM sleep and total sleep (Fig. 2d).

**Fig. 5:**
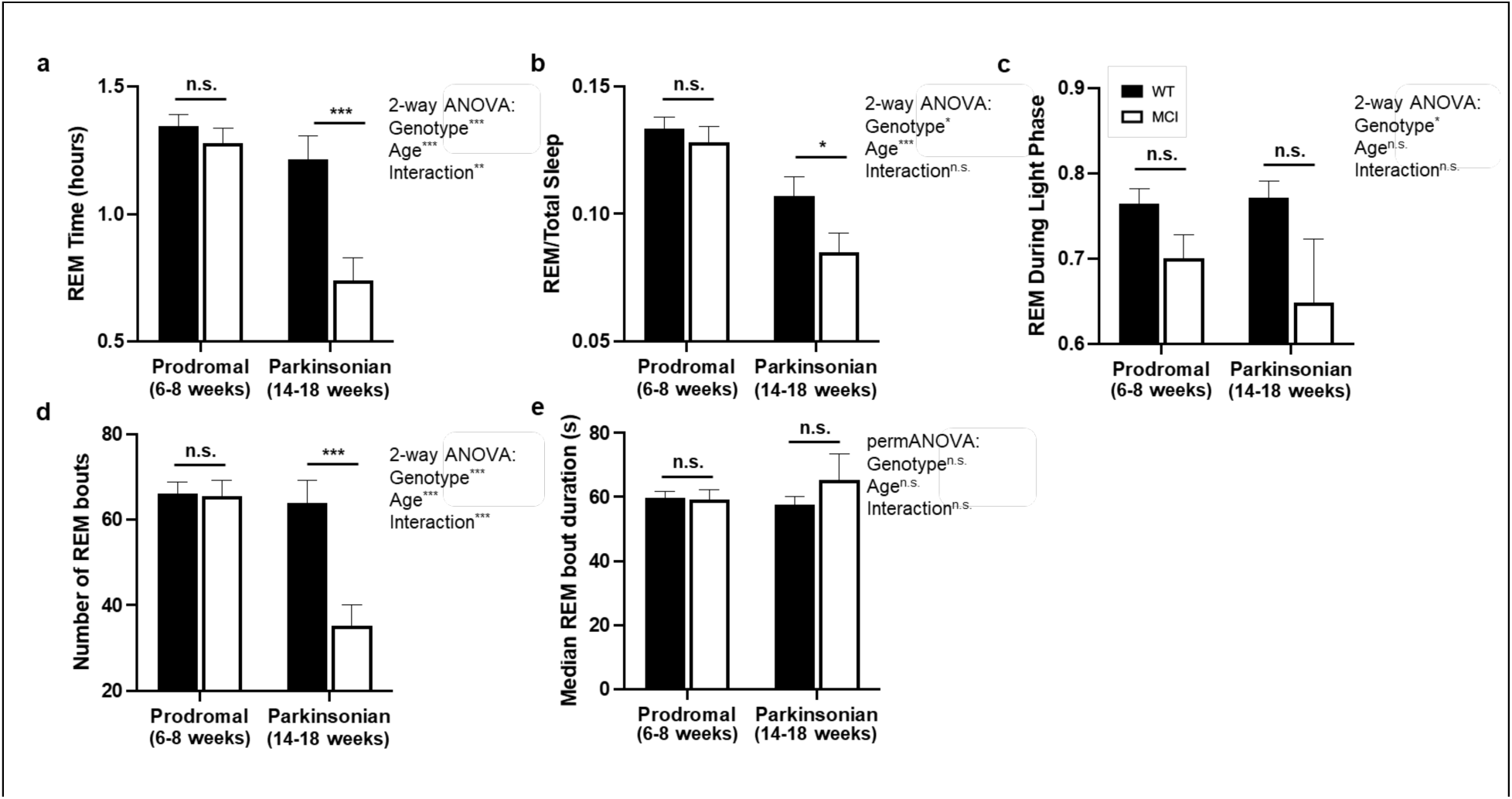
REM sleep amount, distribution, bout number, and median bout duration. **a** – total amount of REM sleep over 24 hours in wildtype (filled bars) and MCI-Park (open bars) mice at 6-8 weeks of age and 14-18 weeks of age. **b** – proportion of total sleep that is REM sleep (REM sleep amount/total sleep amount) in wildtype (filled bars) and MCI-Park (open bars) mice at 6-8 weeks of age (solid bars) and 14-18 weeks of age (open bars). **c** – proportion of REM sleep during the light phase of the light-dark cycle in wildtype (filled bars) and MCI-Park (open bars) mice at 6-8 weeks of age (solid bars) and 14-18 weeks of age (open bars). **d** – number of REM bouts over 24 hours in wildtype (filled bars) and MCI-Park (open bars) mice at 6-8 weeks of age and 14-18 weeks of age. **e** – median duration of REM bouts in wildtype (filled bars) and MCI-Park (open bars) mice at 6-8 weeks of age and 14-18 weeks of age. *, *p*<0.05; **, *p*<0.01; ***, *p*<0.001. *N*=10-24 mice per genotype per age.

There were also significant differences in the number of REM bouts (Fig. 5d) between genotypes (*p*<0.001, F(1,65)=15.75) and age groups (*p*<0.001, F(1,65)=18.15), with MCI-Park and older mice having fewer overall REM bouts compared to wildtype and younger mice, respectively, as reflected by a significant genotype by age interaction (*p*<0.001, F(1,65)=13.00). There were no significant differences in the median REM bout duration (Fig. 5e) between genotypes or ages. This indicates that older MCI-Park mice exhibit fewer bouts of REM sleep without a significant change in bout length.

In conjunction with these reductions in the amount of time spend in REM sleep and the number of REM bouts, significant differences in the inter-REM interval (i.e., duration of time between REM bouts) were also observed between genotypes (*p*<0.001, F(1,65)=15.20) and ages (*p*<0.001, F(1,50)=9.02), with MCI-Park and older mice demonstrating longer intervals between REM bouts. Together, this constellation of REM sleep differences demonstrates a profound dysregulation of REM sleep in MCI-Park mice compared to wildtypes, particularly in the older parkinsonian mice.

### The MCI-Park sleep phenotype was robust

To assess the robustness of the MCI-Park sleep phenotype, an independent study was performed by the Dan laboratory (University of California Berkeley). Sleep was recorded and analyzed as described in the Methods. Although the experimental protocol, sleep recording system, and analysis software were different between the Dan and Turek laboratories, as shown in Supplemental Figure 1, MCI-Park mice in the Dan study exhibited increased wakefulness and decreased NREM sleep, primarily during the light phase, at both the younger (5-8 weeks) and older (14-17 weeks) ages, as observed in the Turek laboratory (Fig. 2). MCI-Park mice in the Dan study also exhibited increased fragmentation, with the older MCI-Park mice experiencing more wake and NREM sleep bouts, as well as decreased NREM bout duration (Supplemental Figure 2), similar to the findings in the Turek laboratory (Fig. 3). In both laboratories, there was a shift towards lower frequencies in the EEG power spectrum (Table 1, Supplemental Figure 3), and REM sleep was impaired, with significantly decreased REM sleep amount, primarily during the light phase, and fewer bouts of REM sleep (Fig. 5, Supplemental Figure 4). The consistency of the findings from separate laboratory environments using different protocols and equipment indicates the robust and reproducible nature of the sleep-wake abnormalities in the MCI-Park model.

## Discussion

The studies described here revealed that the MCI-Park mouse model of PD exhibits dramatically disrupted sleep-wake regulation, characterized by increased wakefulness, decreased NREM sleep, reduced diurnal rhythms of sleep, increased fragmentation, altered EEG patterns, and profoundly impaired REM sleep. Some of these sleep abnormalities manifested prior to the onset of overt motor dysfunction, whereas others were more strongly correlated with PD-like motor disability. Importantly, these sleep disturbances mirror those commonly reported by and observed in PD patients, notably insomnia, excessive daytime sleepiness, increased fragmentation, and changes to sleep architecture^5,6^. Furthermore, the key findings of increased wakefulness, decreased sleep, and increased fragmentation were found in two different laboratory settings, highlighting the robustness of this phenotype. Thus, the MCI-Park mouse provides a unique and powerful model in which PD-related sleep-wake disturbances can be studied.

MCI-Park mice exhibited a significant increase in wakefulness, with corresponding reductions in NREM and REM sleep. Sleep in MCI-Park mice also was more fragmented, which manifested as increased numbers of wake and NREM sleep bouts, decreased REM sleep bouts, more state shifts, and a decreased median NREM bout duration. These patterns mirror the insomnia experienced by PD patients, with difficulty falling asleep, staying asleep, and consolidating sleep without interruption^2–8^. The temporal distribution of sleep was also altered in MCI-Park mice, with decreases in the proportion of NREM sleep and total sleep occurring during the light phase, or “right” time of day for these nocturnal animals. This finding, in conjunction with the increased fragmentation noted above, again mirror the excessive daytime sleepiness and frequent nocturnal awakenings experienced by PD patients^5,8^, which indicate impairment in the overall temporal regulation of sleep.

Analysis of brain wave activity by measuring the EEG power spectral density revealed consistent decreases in relative alpha power across all vigilance states in MCI-Park compared to wildtype mice, as well as a relative increase in delta power during NREM sleep. This complex of changes is consistent with an overall slowing of the EEG, with a shift towards greater power of the slower frequency bands (e.g., delta) and lesser power of the higher frequency bands (e.g., alpha). Such changes have been observed globally in PD patients^11^, and in certain mouse models of PD^14^. Interestingly, decreases in relative alpha power have been proposed as a potential predictor of cognitive impairment in PD^32^, and alterations in alpha power have been linked to cognition and neuropsychology performance^16^, as well as to motivation and incentivized behavior in PD^19^. Future studies of MCI-Park mice may generate hypotheses regarding the role of quantitative EEG analysis as a non-invasive marker for diagnostic, prognostic, and risk-stratification purposes.

Despite the importance of sleep quality to PD patients, the mechanisms responsible for its disruption with disease progression are poorly understood. In large measure, this gap reflects the shortcomings of commonly used PD models. Such models often manifest sleep abnormalities, however the phenotypes are highly variable, inconsistent across different contexts, and typically fail to reproduce the spectrum of abnormalities reported in human PD. For example, studies using the neurotoxin model 1-methyl-4-phenyl-1,2,3,6-tetrahydropyridine (MPTP) are inconsistent across species, with non-human primate models exhibiting sleep-wake abnormalities resembling those in human PD^33^, whereas mice exposed to MPTP have sleep alterations that differ from those observed in patients^6,34^. Mice over-expressing mutant human *α*-synuclein sleep less, experience increased wakefulness, display altered temporal distribution of sleep, and demonstrate a shift in EEG spectral density to lower frequency bands^14,26^; suggesting a PD-like disfunction in sleep. The MitoPark model exhibits increased fragmentation and reduced REM sleep with overall hypersomnia, particularly during the dark (active) phase of the light cycle^15^. Mice lacking VMAT2 (and degeneration of dopaminergic neurons) have a shorter sleep latency defined behaviorally, but not other features of the PD sleep phenotype^33^. Other models have been used to study the response to sleep deprivation^13^, but not the regulation of sleep per se.

In contrast to these other models, the MCI-Park model demonstrates a more complete reproduction of the profound and progressive sleep-wake abnormalities observed in PD patients. There are several differences between the MCI-Park model and other commonly used models. Unlike most other models, the MCI-Park model is based upon an intersectional genetics strategy that selectively targets mitochondrial complex I (MCI) in dopaminergic neurons^29^. An acquired loss of MCI function in dopaminergic neurons is a hallmark of idiopathic PD^35^. Unlike other models, this targeted genetic intervention faithfully recapitulates the progressive, regionally-specific deficits in dopaminergic signaling thought to occur in human PD. As in humans with PD^36^, dopaminergic dysfunction in the MCI-Park model is first evident in the axons innervating the motor regions of the striatum, progressing to regions of the associative or limbic striatum later^29^. This axonal dysfunction is mirrored by a lateral to medial temporal gradient in the dysfunction of dopaminergic neurons in the mesencephalon. This progressive neuropathology is critical to the staging of motor deficits and has provided fundamental new insight into the network mechanisms driving these defining features of PD^29^.

This staging is very likely to provide insight into the network mechanisms underlying sleep deficits as well. For example, several sleep traits, like sleep fragmentation, were disrupted in MCI-Park mice at the earliest time points studied – prior to the onset of parkinsonian motor deficits – pointing to a potential role of striatal dopamine release (as opposed to dopamine release elsewhere) in the ability to sustain NREM sleep bouts. Indeed, there are compelling reasons to think that sleep-wake transitions are strongly influenced by substantia nigra pars reticulata GABAergic neurons whose activity is directly regulated by the striatum^37,38^. Other sleep traits that did not change with age in wildtype mice (e.g., the amount of REM sleep) were progessively impaired in MCI-Park mice, pointing to the potential importance of mesencephalic dopamine release or slowly evolving alterations in brain circuitry triggered by deficits in dopaminergic deficits. Additional studies will be necessary to more clearly define the relationship between dysfunction in specific brain circuits and specific sleep deficits. Again, the staging of pathology in the MCI-Park mice will allow this effort to move forward. The other take-away from our studies is that although many of the sleep deficits seen in PD patients are not responsive to levodopa therapy^39^, this does not mean they are not attributable to the loss of dopaminergic neurons. Levodopa therapy only restores one aspect of dopaminergic signaling – the steady, basal level of dopamine release. The spatiotemporal pattern of dopamine release, which may be critical to the activity of brain circuits controlling sleep architecture, need not be restored by levodopa therapy.

This study has several limitations to be considered. These cross-sectional sleep data provide snapshots of differences at discrete time points. Longitudinal sleep assessment in individual mice over time would provide a better picture of the temporal dynamics of sleep-wake changes, and their relationship with the progression of neuropathology and motor symptoms. In addition, although our studies utilized high-quality EEG/EMG recordings to define vigilance states, they did not incorporate synchronized video recordings, which would allow assessment of RBD-related behaviors – a topic of clear relevance to PD^40^. Finally, although mice are a powerful model in which to study sleep-wake behaviors, there are known species differences in sleep that need to be considered in the interpretation and application of these findings for humans. Despite these limitations, the finding that MCI-Park mice exhibit robust, reproducible, and profound sleep disturbances that are similar in nature to those observed in PD patients is an important advance. Not only does this discovery provide an important insight into the potential mechanisms driving the disruption of sleep in humans with PD, it also provides a new strategy for testing therapeutics that might improve their sleep and quality of life.

## Methods

### Animal Care and Housing

All experimental protocols were reviewed and approved in advance by the Animal Care and Use Committee of Northwestern University and the University of California, Berkeley. All study animals were housed and handled in accordance with Federal Animal Welfare guidelines. Animals used in the Turek laboratory were littermates generated from a breeding colony maintained by the Surmeier laboratory at Northwestern University. Animals used in the Dan laboratory were littermates generated from a breeding colony initially established using breeders from the Surmeier laboratory and then maintained at the University of California, Berkeley. *Dat-cre^-/-^*-*Ndufs2^fl/fl^*mice were used as wildtype control animals and *Dat-cre^+/-^-Ndufs2^fl/fl^*mice (MCI-Park mice; RRID:IMSR_JAX:036313) were used as experimental animals, as previously described^29^. Mice were maintained in constant environmental conditions with a light cycle consisting of 12 hours light followed by 12 hours darkness (LD 12:12) and constant temperature and humidity, as previously described^31^.

The animals were provided *ad libitum* access to a regular chow diet and water. In addition, rodent diets were supplemented with palatable high-calorie energy-dense treats (Turek laboratory: Nutra-Gel Diet, Purified Formula, Bacon Flavor, Product #S5769-TRAY; Bacon Yummies, Product #S05778-1; and Supreme Mini Treats Very Berry Flavor, Product #S05711-1; Bio-Serv, Flemington, NJ; Dan laboratory: ClearH2O DietGel, Product #72-12-5022 and Product #72-10-6000, Westbrook, ME) that were provided fresh each day, as MCI-Park mice were observed to have difficulty maintaining weight at older ages on a standard rodent chow diet alone. Cages were provided with nesting materials and paper strips for environmental enrichment. Mice in the Turek laboratory were group housed until the time of sleep recording, when they were placed in individual sleep recording chambers. Mice in the Dan laboratory were group housed until the time of surgery, after which they were placed in individual cages for recovery and sleep recording. Male and female mice were utilized for all studies.

### Sleep-Wake Recording in the Turek Laboratory

Sleep-wake behavior was recorded as previously described^31^. Mice were implanted with electroencephalography (EEG) and electromyography (EMG) recording electrodes (Pinnacle Technologies, Lawrence, KS) using standard aseptic surgical technique with a stereotaxic apparatus in a dedicated and well-ventilated surgical suite. Prior to surgery, anesthesia was achieved with intraperitoneal (IP) injection of ketamine HCl (98 mg/kg, Vedco Inc., St, Joseph, MO) and xyalizine (10 mg/kg, Akorn Inc, Lake Forest, IL). Surgery consisted of placement of a headmount, containing a plastic 6-pin connector attached to four EEG recording electrodes and two EMG recording electrodes. Four stainless steel screws to anchor the headmounts and serve as EEG electrodes were then screwed into the skull, and the ends of two stainless steel Teflon-coated wires serving as EMG leads were inserted into the nuchal musculature as previously described^41^. The headmount was then attached to the skull using dental acrylic and sutures were used to close the skin incision at the anterior and posterior aspects of the implant. Mice were given a subcutaneous injection of meloxicam (2 mg/kg, Norbrook Laboratories, Newry, Northern Ireland) after surgery and on the following day for analgesia. The mice were allowed to recover from surgery for a minimum of 7 days postoperatively.

After recovery, mice were then acclimated to the cylindrical sleep recording cages (Pinnacle Technologies, Lawrence, KS) within individual acoustically isolated, light tight, and Faraday shielded chambers. The recording cages contained standard corncob bedding and *ad libitum* access to water, regular chow, and palatable dietary supplements. At least 48 hours prior to sleep recording, a tethered preamplifier was plugged into the headmount of each mouse, and then each preamplifier was plugged into an analog-digital converter for collection of EEG/EMG data. EEG/EMG data were collected continuously using Pinnacle Acquisition software (Pinnacle Technologies, Lawrence, KS) and then analyzed. These data were scored in 10 second epochs as wake, NREM sleep, or REM sleep using an automated machine learning-assisted scoring program^30^ supplemented with manual visual inspection.

Epochs unable to be accurately scored were uncommon: no epochs were identified as artifact for any wildtype mice; one older MCI-Park mouse had 6.5% of epochs that were unable to be scored into a particular sleep-wake state, so these were labeled as artifact and the mouse was excluded from further analysis; one younger MCI-Park mouse had 0.2% of epochs scored as artifact; and another older MCI-Park had 0.6% of epochs scored as artifact. One older MCI-Park mouse had poor quality recordings from one of the two EEG channels used for data collection (i.e., EEG1). The vigilance states (wake, NREM sleep, REM sleep) for this animal were adequately identified using the other EEG channel (i.e., EEG2), so the animal was included in the analysis of sleep-wake states and fragmentation traits. However, it was excluded from relative EEG power calculations and analyses, which were performed using recordings from EEG1 on all other mice.

### Sleep-Wake Recording in the Dan Laboratory

For surgical procedures, mice were anesthetized with 1.5% isoflurane and placed in a stereotaxic frame. Body temperature was maintained using a heating pad. The skin was incised to expose the skull after asepsis, and connective tissue was removed. To implant EEG and EMG recording electrodes, two stainless steel screws were inserted into the skull 2.5 mm from the midline and 3 mm posterior to the bregma, and two EMG electrodes were also inserted into the neck muscles. A reference screw for grounding was placed on top of the left cerebellum.

Insulated leads from the EEG and EMG electrodes were soldered to a pin header, which was secured to the skull using dental cement.

After a minimum of 7 days of undisturbed post-surgery recovery, behavioral experiments were carried out in home cages placed in sound-attenuating boxes. EEG and EMG electrodes were connected to flexible recording cables via a mini-connector. Recordings started after 1-2 days of habituation and continued for 3-4 days on a 12-h dark/12-h light cycle. EEG and EMG signals were recorded with a TDT RZ10x/PZ5-32 or LR10-SK1 system. Spectral analysis was carried out using fast Fourier transform (FFT), and brain states were scored in 5 second epochs and classified into wake (desynchronized EEG and high EMG activity), NREM sleep (synchronized EEG with high delta power (1-4 Hz) and low EMG activity) and REM sleep (high EEG theta power (6-9 Hz), low EEG delta power and low EMG activity). The classification of brain states (wake, NREM or REM) was completed using a custom-written graphical user interface (‘AccuSleep’ GitHub package; DOI:10.5281/zenodo.10055607)^42^, with automated scoring followed by manual correction.

### Statistical Analysis

For results obtained in the Turek laboratory, sleep measures were examined by three-way Analysis of Variance (ANOVA) to identify genotype (wildtype *vs* MCI-Park), age (prodromal/6-8 weeks of age *vs* parkinsonian/14-18 weeks of age), sex (male *vs* female), or interaction effects. Three-way ANOVA revealed no significant effects of sex for any of the sleep measures analyzed, so male and female mice were combined for subsequent analyses using two-way ANOVA to evaluate for effects of genotype, age, and genotype-by-age interactions. For measures that satisfied normality tests, a Generalized Linear Models (GLM) ANOVA with

Tukey-Kramer post-hoc pairwise comparisons and alpha ≤ 0.05 was done using Number Crunchers Statistical Software (NCSS; https://www.ncss.com/) 2019 (Kayesville, UT) and GraphPad Prism (v10.0.0; GraphPad Software, Boston, MA; RRID:SCR_002798). Bout numbers of vigilance states (wake, NREM sleep, REM sleep) were square-root transformed to achieve normality before analysis. The number of brief arousals and state shifts were log (base 10) transformed to achieve normality before analysis.

Because the variances for median vigilance state bout durations (wake, NREM sleep, REM sleep) were not normal, and were unable to transformed to achieve normality, a non-parametric permutation-based ANOVA (PermANOVA) was employed to evaluate genotype (wildtype *vs* MCI-Park), age (prodromal/6-8 weeks of age *vs* parkinsonian/14-18 weeks of age), or genotype-by-age interaction effects for these measures. 5000 permutations were used with 1000 repetitions bootstrap post-hoc tests (DOI:10.5281/zenodo.10079840). Post-hoc *p*-values were false discovery rate (FDR) corrected. The PermANOVA was performed using the Permuco package (RRID:SCR_022341)^43^ in R Statistical Software (4.3.1, R Core Team, Vienna, Austria, 2023; RRID:SCR_001905). Bootstrap post-hoc analyses were performed using the Car package (RRID:SCR_022137)^44^. Figures were generated using GraphPad Prism (v10.0.0; GraphPad Software, Boston, MA; RRID:SCR_002798).

For results obtained in the Dan laboratory, sleep measures were compared between similarly aged wildtype and MCI-Park mice using t-tests.

## Data Availability

The datasets on sleep-wake traits in MCI-Park and littermate wildtype mice generated in this study are publically available (Northwestern University, DOI:10.5281/zenodo.10079840; University of California Berkeley, DOI: 10.5281/zenodo.10046587). In addition, specific aspects of the data will be made available by the corresponding author (KCS, Northwestern University data) or the co-first author (XH, University of California Berkeley data) in response to reasonable requests from qualified researchers (i.e., researchers affiliated with a university, academic medical center, research institute, or similar institution), pending approval of all senior authors (MHV, YD, DJS, FWT).

## Author Contributions

MHV, YD, DJS, and FWT designed the experiments and obtained funding. KCS, PJ, and MHV performed experiments in the Turek laboratory. XH performed experiments in the Dan laboratory. PGR and DJS developed and provided the mouse model used in the experiments.

KCS, PGR, XH, XL, and MHV performed statistical analyses. All authors assisted with the interpretation of the data. KCS wrote the initial draft of the manuscript, with input from all authors on subsequent revised drafts. All authors read and approved the final version of the manuscript for submission.

## Acknowledgements

This study was funded by the Department of Defense grants W81XWH2110582 (FWT) and W81WH2110749 (DJS), Aligning Science Against Parkinson’s (ASAP) grant 020551 (DJS, YD), and the JPB Foundation (DJS). The funders performed no role in study design, data collection, analysis and interpretation of data, or in writing of this manuscript. We would like to thank J. Sun, R. Xiang, C. Chen, and Y. Liu from the Dan laboratory for assistance in completing the studies at University of California Berkeley described in this manuscript.

## Supplemental Material

**Supplemental Fig. 1:**
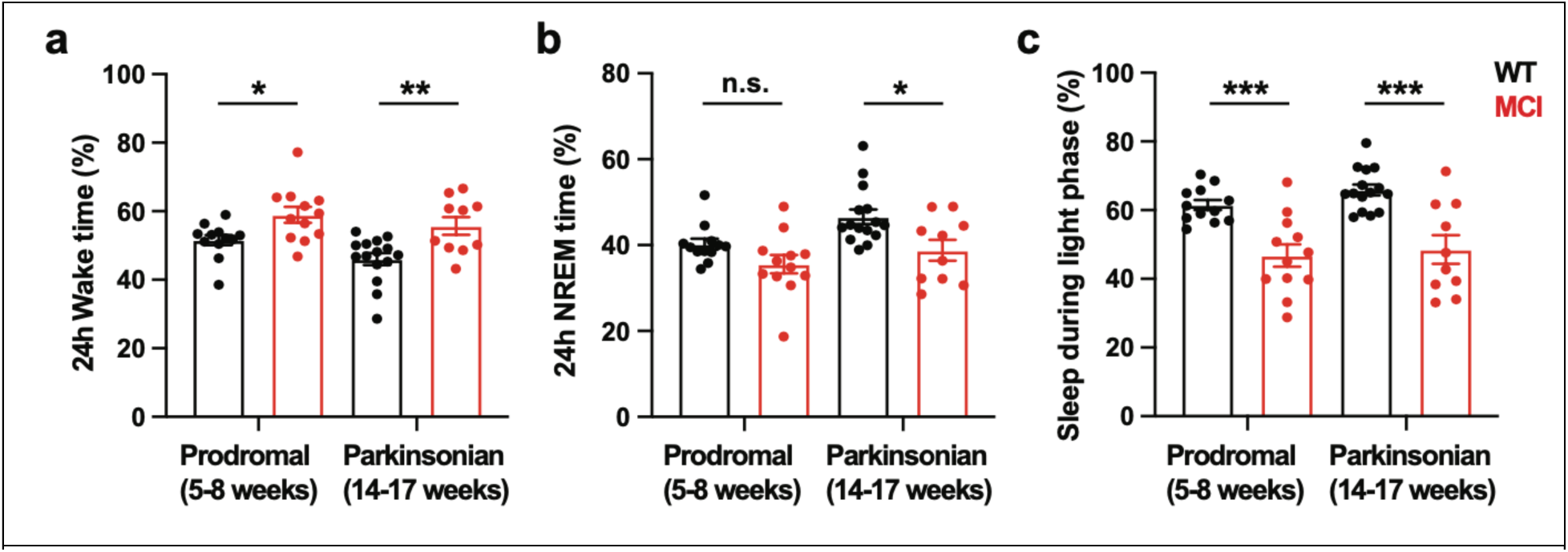
Percentage of time spent in wake (a) and NREM sleep (b), and proportion of sleep during the light phase (c) in wildtype (black) and MCI-Park (red) mice at 5- 8 weeks of age (left panels) and 14-17 weeks of age (right panels). * p<0.05, ** p<0.01, *** p<0.001; t-test.

**Supplemental Fig. 2.**
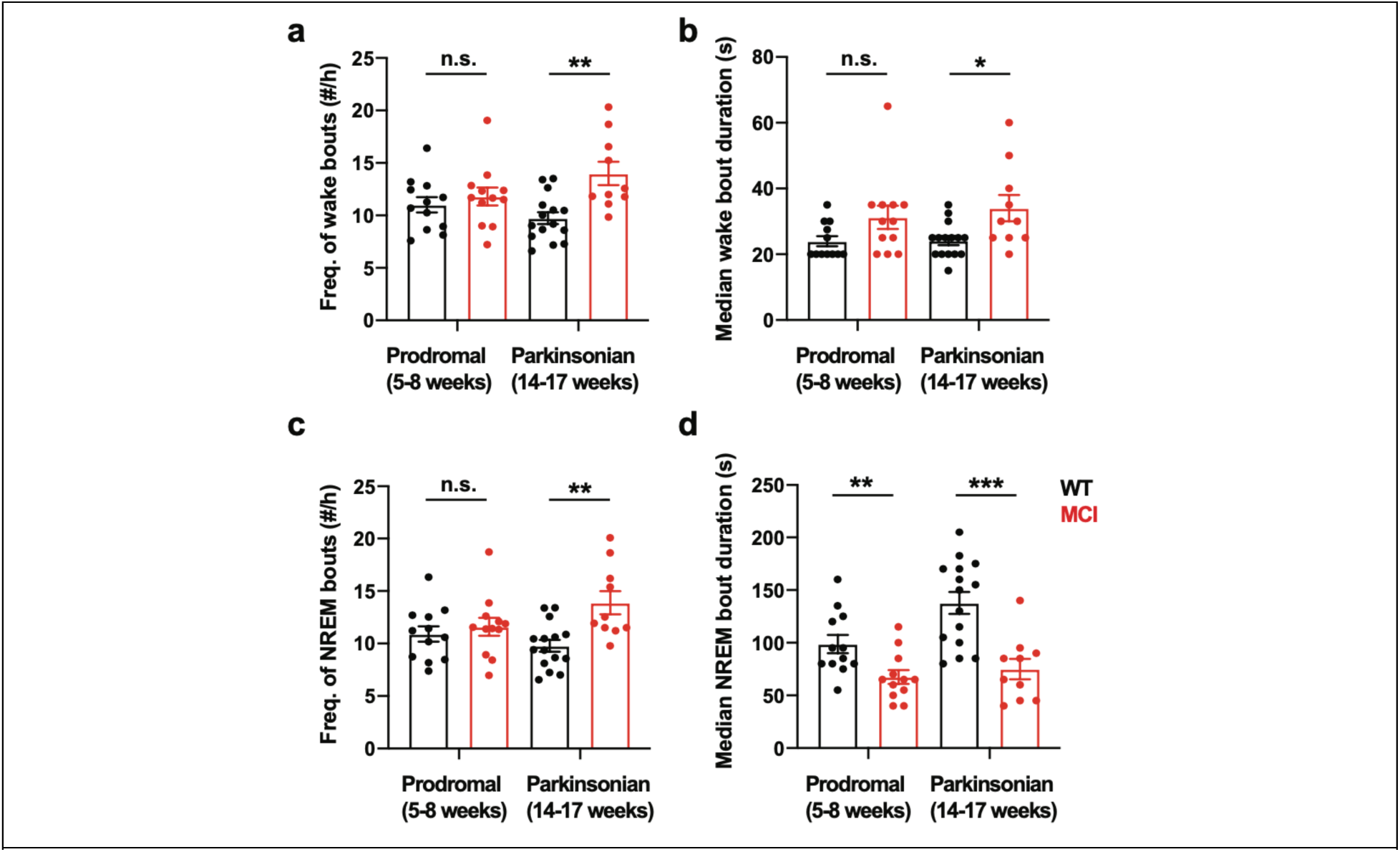
Frequency (# bouts/hr) and median duration (s) of bouts of wake (a, b) and NREM sleep (c, d) in wildtype (black) and MCI-Park (red) mice at 5-8 weeks of age (left panels) and 14-17 weeks of age (right panels). * p<0.05, ** p<0.01, *** p<0.001; t-test.

**Supplemental Fig. 3.**
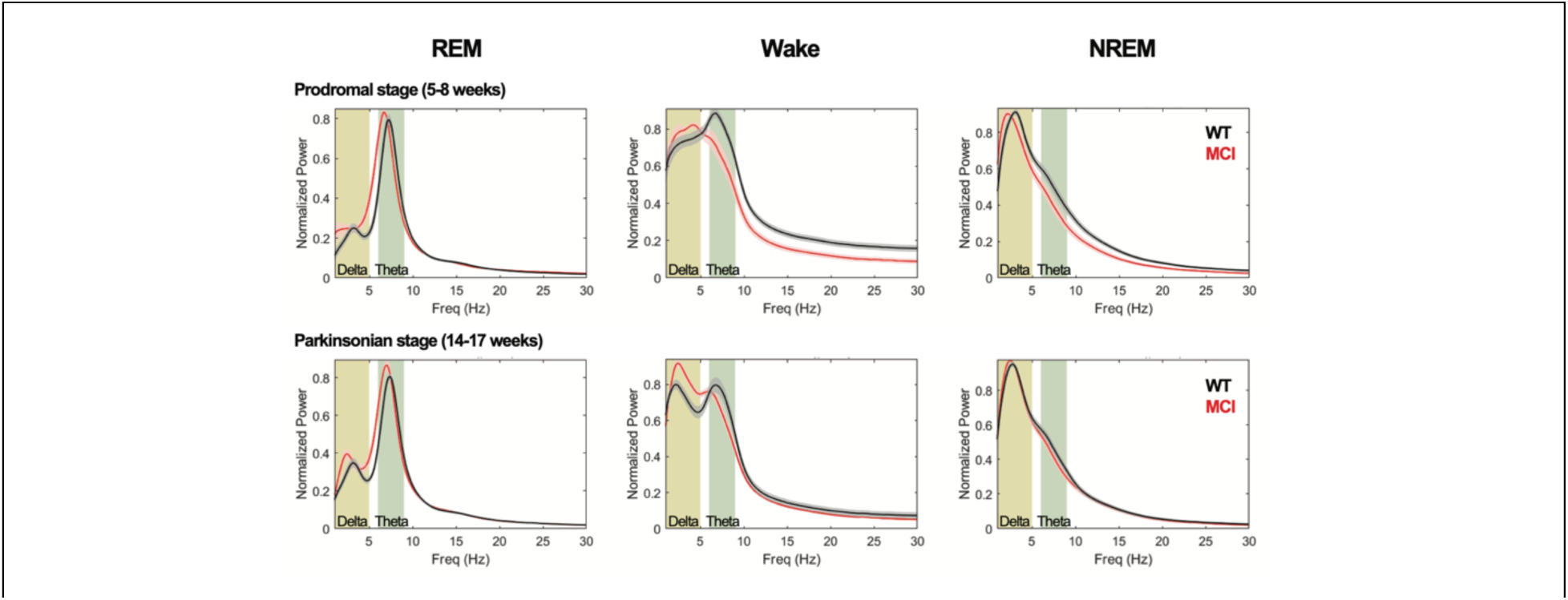
EEG power spectral plots for REM sleep (left panel), wake (middle panel), and NREM sleep (right panel) in wildtype (black) and MCI-Park (red) mice at 5-8 weeks of age (top row) and 14-17 weeks of age (bottom row).

**Supplemental Fig. 4.**
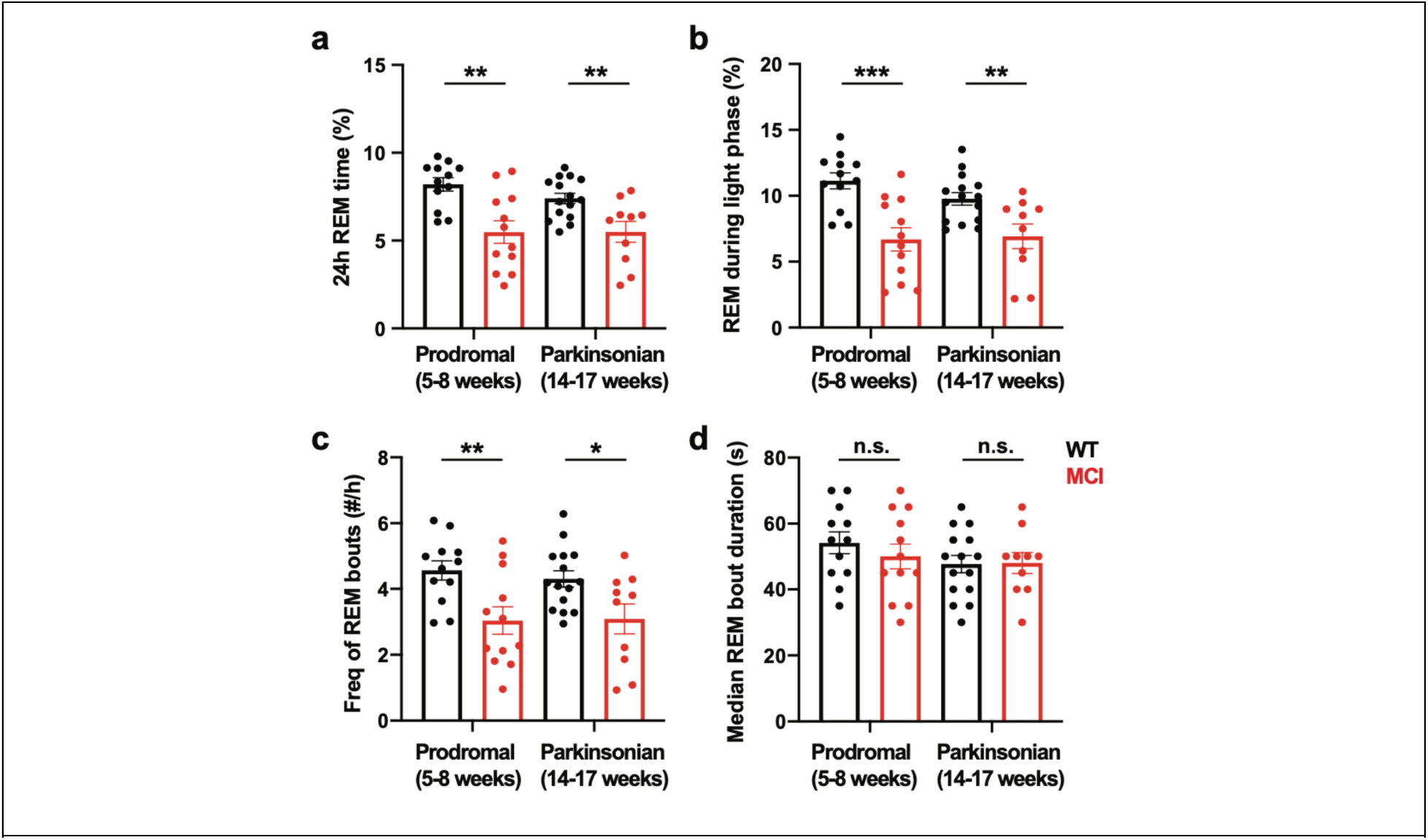
Percentage of time spent in REM sleep (a), proportion of REM sleep during the light phase (b), frequency of REM bouts (# bouts/hr), and median duration of REM bouts (s) in wildtype (black) and MCI-Park (red) mice at 5-8 weeks of age (left panels) and 14- 17 weeks of age (right panels). * p<0.05, ** p<0.01, *** p<0.001; t-test.

**Supplemental Table 1.** Relative EEG spectral power for wake, NREM sleep, and REM sleep, in wildtype (WT) and MCI-Park (MCI) mice at younger (prodromal, age 6-8 weeks) and older (parkinsonian, age 14-18 weeks), calculated as the power in that band divided by total power in that state and expressed as a percentage ± standard error. *N*=10-23 mice per genotype per age.

| State | Power Band | WT Prodromal<br>(6-8 weeks) | WT Parkinsonian<br>(14-18 weeks) | MCI Prodromal<br>(6-8 weeks) | MCI Parkinsonian<br>(14-18 weeks) |
|---|---|---|---|---|---|
| Wake | Delta<br>(0.5-4 Hz) | 30.3±1.7% | 26.0±1.0% | 29.4±1.7% | 31.5±2.5% |
| Wake | Theta<br>(4-8 Hz) | 34.8±0.7% | 39.2±1.1% | 36.5±0.9% | 35.9±1.4% |
| Wake | Alpha<br>(8-11 Hz) | 15.6±0.8% | 15.5±0.5% | 14.1±0.7% | 12.2±1.3% |
| Wake | Sigma<br>(11-15 Hz) | 7.0±0.2% | 7.1±0.2% | 7.2±0.2% | 7.1±0.4% |
| Wake | Beta<br>(15-30 Hz) | 12.2±0.5% | 12.3±0.7% | 12.9±0.4% | 13.3±1.1% |
| NREM | Delta<br>(0.5-4 Hz) | 45.6±1.6% | 37.1±1.8% | 47.3±1.5% | 43.2±1.4% |
| NREM | Theta<br>(4-8 Hz) | 30.5±0.7% | 37.0±0.8% | 30.7±0.8% | 33.7±1.1% |
| NREM | Alpha<br>(8-11 Hz) | 10.3±0.5% | 11.9±0.5% | 9.6±0.4% | 9.9±0.4% |
| NREM | Sigma<br>(11-15 Hz) | 6.7±0.3% | 6.8±0.4% | 6.1±0.2% | 6.0±0.2% |
| NREM | Beta<br>(15-30 Hz) | 6.8±0.3% | 7.2±0.4% | 6.4±0.2% | 7.2±0.4% |
| REM | Delta<br>(0.5-4 Hz) | 22.3±1.3% | 20.4±1.5% | 21.7±1.1% | 23.2±2.5% |
| REM | Theta<br>(4-8 Hz) | 40.5±0.7% | 39.2±1.2% | 41.5±1.0% | 41.0±2.7% |
| REM | Alpha<br>(8-11 Hz) | 16.7±0.7% | 19.6±0.9% | 15.1±0.6% | 14.4±1.5% |
| REM | Sigma<br>(11-15 Hz) | 7.6±0.2% | 7.8±0.4% | 8.2±0.2% | 7.8±0.3% |
| REM | Beta<br>(15-30 Hz) | 12.8±0.3% | 13.0±0.7% | 13.5±0.6% | 13.5±1.0% |

### Sleep Trait Definitions

**%[state]:** % of epochs in that particular state. %wake + %nrem + %rem + %artifact = 100.

%nrem + %rem = %sleep. If an epoch’s score is ’unscored’ then it will be treated as an artifact. [state]/sleep : %[state]/%sleep. nrem/sleep + rem/sleep = 1

**# [state] bouts:** count of the number of bouts of that state

**[state] median bout duration:** median bout length of that particular state in seconds

**brief arousals:** the number of single wake epochs in the middle of a sleep bout

**state shifts:** of the 3 vigilance states (i.e., wake, NREM, or REM), the number of times that the current epoch is not the same as the previous one

**[state] [band] total:** the average raw value of that band (i.e., delta, theta, alpha, sigma, beta) in that particular state. Excluded epochs are excluded from this calculation.

**[state] %[band]:** for each epoch in the period of that particular state, a relative power score is determined and expressed as a percentage. That is, if total power = Pt = raw power delta + raw power theta + raw power alpha + raw power sigma + raw power beta, then the relative power of the band = (raw [band]/Pt)*100. The output given is the mean of the relative[band] values.

Excluded epochs are excluded from this calculation.

## References

1 Poewe, W. et al. Parkinson disease. Nat Rev Dis Primers 3, 17013 (2017). 10.1038/nrdp.2017.13

2 Stefani, A. & Hogl, B. Sleep in Parkinson’s disease. Neuropsychopharmacology 45, 121–128 (2020). 10.1038/s41386-019-0448-y

3 Chahine, L. M., Amara, A. W. & Videnovic, A. A systematic review of the literature on disorders of sleep and wakefulness in Parkinson’s disease from 2005 to 2015. Sleep Med Rev 35, 33–50 (2017). 10.1016/j.smrv.2016.08.001

4 French, I. T. & Muthusamy, K. A. A Review of Sleep and Its Disorders in Patients with Parkinson’s Disease in Relation to Various Brain Structures. Front Aging Neurosci 8, 114 (2016). 10.3389/fnagi.2016.00114

5 Gros, P. & Videnovic, A. Sleep and Circadian Rhythm Disorders in Parkinson’s Disease. Curr Sleep Med Rep 3, 222–234 (2017). 10.1007/s40675-017-0079-y

6 Hunt, J. et al. Sleep and circadian rhythms in Parkinson’s disease and preclinical models. Mol Neurodegener 17, 2 (2022). 10.1186/s13024-021-00504-w

7 Maggi, G., Vitale, C., Cerciello, F. & Santangelo, G. Sleep and wakefulness disturbances in Parkinson’s disease: A meta-analysis on prevalence and clinical aspects of REM sleep behavior disorder, excessive daytime sleepiness and insomnia. Sleep Med Rev 68, 101759 (2023). 10.1016/j.smrv.2023.101759

8 Mantovani, S., Smith, S. S., Gordon, R. & O’Sullivan, J. D. An overview of sleep and circadian dysfunction in Parkinson’s disease. J Sleep Res 27, e12673 (2018). 10.1111/jsr.12673

9. 9 Malhotra, R. K. Neurodegenerative Disorders and Sleep. Sleep Med Clin 13, 63–70 (2018). 10.1016/j.jsmc.2017.09.006

10 Morita, A., Kamei, S. & Mizutani, T. Relationship between slowing of the EEG and cognitive impairment in Parkinson disease. J Clin Neurophysiol 28, 384–387 (2011). 10.1097/WNP.0b013e3182273211

11 Soikkeli, R., Partanen, J., Soininen, H., Paakkonen, A. & Riekkinen, P., Sr. Slowing of EEG in Parkinson’s disease. Electroencephalogr Clin Neurophysiol 79, 159–165 (1991). 10.1016/0013-4694(91)90134-p

12 Dushanova, J., Philipova, D. & Nikolova, G. Beta and gamma frequency-range abnormalities in parkinsonian patients under cognitive sensorimotor task. J Neurol Sci 293, 51–58 (2010). 10.1016/j.jns.2010.03.008

13 Liu, X. et al. Altered Motor Performance, Sleep EEG, and Parkinson’s Disease Pathology Induced by Chronic Sleep Deprivation in Lrrk2(G2019S) Mice. Neurosci Bull 38, 1170–1182 (2022). 10.1007/s12264-022-00881-2

14 McDowell, K. A., Shin, D., Roos, K. P. & Chesselet, M. F. Sleep dysfunction and EEG alterations in mice overexpressing alpha-synuclein. J Parkinsons Dis 4, 531–539 (2014). 10.3233/JPD-140374

15 Fifel, K., Yanagisawa, M. & Deboer, T. Mechanisms of Sleep/Wake Regulation under Hypodopaminergic State: Insights from MitoPark Mouse Model of Parkinson’s Disease. Adv Sci (Weinh) 10, e2203170 (2023). 10.1002/advs.202203170

16 Jaramillo-Jimenez, A. et al. Resting-state EEG alpha/theta ratio related to neuropsychological test performance in Parkinson’s Disease. Clin Neurophysiol 132, 756–764 (2021). 10.1016/j.clinph.2021.01.001

17 Klimesch, W. EEG alpha and theta oscillations reflect cognitive and memory performance: a review and analysis. Brain Res Brain Res Rev 29, 169–195 (1999). 10.1016/s0165-0173(98)00056-3

18 Ye, Z., Heldmann, M., Herrmann, L., Bruggemann, N. & Munte, T. F. Altered alpha and theta oscillations correlate with sequential working memory in Parkinson’s disease. Brain Commun 4, fcac096 (2022). 10.1093/braincomms/fcac096

19 Zhu, M. et al. Altered EEG alpha and theta oscillations characterize apathy in Parkinson’s disease during incentivized movement. Neuroimage Clin 23, 101922 (2019). 10.1016/j.nicl.2019.101922

20 Dauvilliers, Y. et al. REM sleep behaviour disorder. Nat Rev Dis Primers 4, 19 (2018). 10.1038/s41572-018-0016-5

21 Hogl, B., Stefani, A. & Videnovic, A. Idiopathic REM sleep behaviour disorder and neurodegeneration - an update. Nat Rev Neurol 14, 40–55 (2018). 10.1038/nrneurol.2017.157

22 Van Den Berge, N. & Ulusoy, A. Animal models of brain-first and body-first Parkinson’s disease. Neurobiol Dis 163, 105599 (2022). 10.1016/j.nbd.2021.105599

23 Taylor, T. N., Caudle, W. M. & Miller, G. W. VMAT2-Deficient Mice Display Nigral and Extranigral Pathology and Motor and Nonmotor Symptoms of Parkinson’s Disease. Parkinsons Dis 2011, 124165 (2011). 10.4061/2011/124165

24 Shen, Y. et al. Propagated alpha-synucleinopathy recapitulates REM sleep behaviour disorder followed by parkinsonian phenotypes in mice. Brain 143, 3374–3392 (2020). 10.1093/brain/awaa283

25 Taguchi, T. et al. alpha-Synuclein BAC transgenic mice exhibit RBD-like behaviour and hyposmia: a prodromal Parkinson’s disease model. Brain 143, 249–265 (2020). 10.1093/brain/awz380

26 Rothman, S. M. et al. Neuronal expression of familial Parkinson’s disease A53T alpha-synuclein causes early motor impairment, reduced anxiety and potential sleep disturbances in mice. J Parkinsons Dis 3, 215–229 (2013). 10.3233/JPD-120130

27 Kudo, T., Loh, D. H., Truong, D., Wu, Y. & Colwell, C. S. Circadian dysfunction in a mouse model of Parkinson’s disease. Exp Neurol 232, 66–75 (2011). 10.1016/j.expneurol.2011.08.003

28 Morawska, M. M. et al. Slow-wave sleep affects synucleinopathy and regulates proteostatic processes in mouse models of Parkinson’s disease. Sci Transl Med 13, eabe7099 (2021). 10.1126/scitranslmed.abe7099

29 Gonzalez-Rodriguez, P. et al. Disruption of mitochondrial complex I induces progressive parkinsonism. Nature 599, 650–656 (2021). 10.1038/s41586-021-04059-0

30 Gao, V., Turek, F. & Vitaterna, M. Multiple classifier systems for automatic sleep scoring in mice. J Neurosci Methods 264, 33–39 (2016). 10.1016/j.jneumeth.2016.02.016

31 Winrow, C. J. et al. Uncovering the genetic landscape for multiple sleep-wake traits. PLoS One 4, e5161 (2009). 10.1371/journal.pone.0005161

32 Guo, Y. et al. Predictors of cognitive impairment in Parkinson’s disease: a systematic review and meta-analysis of prospective cohort studies. J Neurol 268, 2713–2722 (2021). 10.1007/s00415-020-09757-9

33 Davin, A. et al. Early onset of sleep/wake disturbances in a progressive macaque model of Parkinson’s disease. Sci Rep 12, 17499 (2022). 10.1038/s41598-022-22381-z

34 Monaca, C. et al. Vigilance states in a parkinsonian model, the MPTP mouse. Eur J Neurosci 20, 2474–2478 (2004). 10.1111/j.1460-9568.2004.03694.x

35 Surmeier, D. J., Obeso, J. A. & Halliday, G. M. Selective neuronal vulnerability in Parkinson disease. Nat Rev Neurosci 18, 101–113 (2017). 10.1038/nrn.2016.178

36 Kordower, J. H. & Burke, R. E. Disease Modification for Parkinson’s Disease: Axonal Regeneration and Trophic Factors. Mov Disord 33, 678–683 (2018). 10.1002/mds.27383

37 Lai, Y. Y., Kodama, T., Hsieh, K. C., Nguyen, D. & Siegel, J. M. Substantia nigra pars reticulata-mediated sleep and motor activity regulation. Sleep 44 (2021). 10.1093/sleep/zsaa151

38 Liu, D. et al. A common hub for sleep and motor control in the substantia nigra. Science 367, 440–445 (2020). 10.1126/science.aaz0956

39 Seppi, K. et al. The Movement Disorder Society Evidence-Based Medicine Review Update: Treatments for the non-motor symptoms of Parkinson’s disease. Mov Disord 26 **Suppl 3**, S42–80 (2011). 10.1002/mds.23884

40 Bohnen, N. I. & Hu, M. T. M. Sleep Disturbance as Potential Risk and Progression Factor for Parkinson’s Disease. J Parkinsons Dis 9, 603–614 (2019). 10.3233/JPD-191627

41 Bowers, S. J. et al. Repeated sleep disruption in mice leads to persistent shifts in the fecal microbiome and metabolome. PLoS One 15, e0229001 (2020). 10.1371/journal.pone.0229001

42 Barger, Z., Frye, C. G., Liu, D., Dan, Y. & Bouchard, K. E. Robust, automated sleep scoring by a compact neural network with distributional shift correction. PLoS One 14, e0224642 (2019). 10.1371/journal.pone.0224642

43 Frossard, J. & Renaud, O. Permutation Tests for Regression, ANOVA, and Comparison of Signals: The permuco Package. Journal of Statistical Software 99, 1–32 (2021). 10.18637/jss.v099.i15

44. Fox, J., Wesiberg, S. An R Companion to Applied Regression. Third edn, (Sage, 2019).

